# Grasp context-dependent uncertainty alters the relative contribution of anticipatory and feedback-based mechanisms in object manipulation

**DOI:** 10.1101/2024.01.24.577057

**Authors:** Swarnab Dutta, Varadhan SKM

## Abstract

Predictive control within dexterous object manipulation while allowing for the choice of contact points has been shown to employ a predominantly feedback-based force modulation. Anticipation is thought to be facilitated through the internal representation of the object dynamics being integrated and updated on a trial-to-trial basis with the feedback of contact locations on the object. This is as opposed to the classically studied memory representation-based fingertip force control for grasping with pre-selected contact locations. We designed a study to examine this grasp context-dependent asymmetry in sensorimotor integration by introducing binary uncertainty about the grasp type before movement initiation within the framework of motor planning. An inverted T-shaped instrumented object was presented to 24 participants as the manipulandum, and they were asked to reach, grasp, and lift it while minimising the roll. We dissociated the planning and the execution phases by pseudo-randomly manipulating the availability of visual contact cues on the object after movement onset. Our results suggest that uncertainty about the grasp type during movement preparation anterogradely modulated the differential weighting of feedback and feedforward mechanisms in anticipatory coordination.

## Introduction

Anticipatory mechanisms underpin the natural ability of humans to manipulate objects, as a sensory delay would be involved with an entirely feedback-based sensorimotor control. These processes dynamically adjust the motor commands to the physical properties of the concerned object depending on the intent of the interaction (Johansson, 2007; Johansson & Flanagan, 2009). Systematically varying the object dynamics, such as eccentric loading, has been considered to add an element of dexterity in object manipulation tasks besides scaling the fingertip forces to the object weight. The desired behavioural outcome of preventing object roll while lifting (akin to preventing spillage while drinking from a cup of hot coffee) consequently necessitates the production of compensatory moments of force on the object at lift onset in an anticipatory manner (J. Lukos et al., 2007; J. R. Lukos et al., 2008; Salimi et al., 2000, 2003).

Control of fingertip forces in object manipulation has typically been studied through the constrained grasp paradigm wherein the objects are grasped at visually indicated points of contact. These learnt manipulations over consecutive trials have been proposed to involve stored information from associations between previous hand-object experiences and the object properties, also termed *sensorimotor memory* (A. M. Gordon et al., 1993; Johansson & Cole, 1992; Johansson & Westling, 1988; Westling & Johansson, 1984). Consider that the resultant forces and moments on the object could be altered by varying the point of application of the fingertip forces besides modulating the fingertip forces. The unconstrained grip paradigm, wherein free choice of finger placement on the object is offered, has thus been pioneering in enabling the understanding of the somewhat overlooked coordination between fingertip position and the forces while lending greater ecological validity to manipulation tasks. Upon being presented with the choice of contact points in an asymmetrically loaded object, the anticipation of the external moment on the object, vis-a-vis the task goal, was accomplished through precise coordination between the fingertip forces and finger placement. The variability here in fingertip placement notwithstanding, which results from the absence of visual cues to guide grasping, skilled manipulation within the unconstrained paradigm can be accurately performed because participants modulate fingertip forces as a function of the fingertip locations on a trial-by-trial basis.

Going beyond the classically established memory representation-based fingertip force control, a predominantly online feedback of fingertip position-based mechanism has been thought to drive the change in the force distribution every time the object is grasped at novel contact points (Fu et al., 2010; Fu & Santello, 2014; Mojtahedi et al., 2015). This theoretical framework suggests a greater reliance on the sensory feedback of fingertip placement locations in the unconstrained condition in that the participants integrate information about the fingertip position with the sensorimotor memory of forces from previous manipulations to build a robust representation of the object dynamics. The presence of greater high-frequency feedback-based corrections and non-bell-shaped grip force rate (GFR) profiles during the loading phase in unconstrained grasping has later lent more support to this line of reasoning (Mojtahedi et al., 2015). Non-invasive brain stimulation studies in the subsequent years have extended this line of research by showing that allowing or preventing choice of contact points modulates distinct areas in the brain as well as the corticospinal excitability (CSE) at object contact (Davare et al., 2019; Marneweck et al., 2018; Marneweck & Grafton, 2020). *Parikh et al.* have probed this further and found that depending on the grasp context, the relative contribution of memory and online feedback differentially engages the primary motor (M1) and the somatosensory (S1) cortices (Parikh et al., 2020).

To our knowledge, none of these studies have investigated the role of a binary uncertainty regarding the grasp context during the pre-movement planning phase, i.e., the reaction time before a voluntary movement is initiated (Wong et al., 2015). In this light, we attempted to examine the consequence of a binary uncertainty about the grasp context during movement preparation in sensorimotor integration. For this, we introduced a systematic dissociation between the planning and the execution phase for the aforementioned adaptation task, i.e., lifting an eccentrically loaded object without tilting.

The role of motor planning when there is ambiguity about the goal information has been investigated by introducing uncertainty through controlled experiments in many different ways: by combining potential goals with interferences (Arai et al., 2004; Walker et al., 1997), deferring goal information (Chapman et al., 2010; Ghez et al., 1997), presenting noisy visual cues (Hudson et al., 2007; Resulaj et al., 2009), or demanding high-level cognitive decision-making during movement (Song & Nakayama, 2009). For the current study, we introduced a binary uncertainty about the grasp context during the movement preparation phase. Specifically, on certain *catch* trials within a regular constrained block, unbeknownst to the participants, the target cues for finger placement were extinguished at movement onset, essentially transforming from a constrained grasp to an unconstrained grasp (see *methods and materials* for details). Such extinguishing of the visual cues during reaching has been employed in tasks that involved reaching towards a visual target [for review, see (Sarlegna & Mutha, 2015)]. However, those studies only focused on the kinematics of reaching and did not involve any object manipulation. The role of the *catch* trials was to dissociate the information in the planning phase from the information in the execution phase. This, in turn, was supposed to induce uncertainty in the participants about the grasp context during movement preparation. Note that the uncertainty in our study is different from the sensorimotor uncertainty intrinsic to the unconstrained grasp condition (Davare et al., 2019). The rest of the trials within this block, called the *test* trials, had visual feedback of contact cues available throughout the task. It is important to highlight that the constrained trials were essentially identical to the *test* trials after the uncertainty was resolved at movement onset in the latter.

In literature, motor planning has been shown to modulate the preparatory states of the primary motor cortex (M1) (Crammond & Kalaska, 2000; Tanji & Evarts, 1976). More recently, the primary somatosensory cortex (S1) that has been causally implicated in unconstrained grasping along with M1 (Parikh et al., 2020), has also been shown to elicit signatures of activation before movement onset during object manipulation (Ariani et al., 2022; Gale et al., 2021). These results, together with the finding that freedom of choice of finger placement as opposed to restrictive contact cues influences brain dynamics differentially during the planning phase (McGurrin, 2017), led us to hypothesise the following:

H1: Uncertainty before movement initiation about the presence of visual grasp targets at contact would elicit kinetic signatures during the loading phase.

## Methods and Materials

### Participants

A total of 24 individuals (12 females, 12 males, mean age: 25.6 ± 3.4 years) with normal or corrected-to-normal vision, all right-handed (self-reported), volunteered to participate in the study. None of the participants had any history of neurological or musculoskeletal impairments of the involved upper limb. All participants were naive to the experimental purpose of the study and provided written informed consent to participate in the experiment. The experimental procedures were approved by the Institutional Ethics Committee of the Indian Institute of Technology Madras (IEC/2021-03/SKM/14), and the experimental sessions were conducted in strict adherence to the approved procedures. The participants were compensated financially according to the institute norms for partaking in the study, which lasted ∼1 hour.

### Experimental setup

Two six-dimensional force and moment transducers (Nano 17, Force resolution: Tangential: 0.0125 N, Normal: 0.0125 N, ATI Industrial Automation, Garner, NC, USA) were instrumented into an inverted T-shaped manipulandum made of acrylic to record the forces and moments exerted from the index and the thumb fingertips (Fig. 1). To create the grasp surfaces, two panels made of polycarbonate and covered with sandpaper (3M, P320) were mounted vertically on each sensor. The manipulandum measured *grip* and *load* forces (normal and tangential to the graspable surfaces, respectively) and each effector’s *centre of pressure* (COP). The graspable panels had a longitudinally flanged portion to impede the panel’s bending from the normal forces. A box made with acrylic with three compartments (middle and one each on either side of the centre) was added at the base of the manipulandum for external weights to change the mass distribution of the assembly, which weighed 410 g as a whole without the external weight. An inertial measurement unit (IMU: Resolution: 16bits, Range: 2000°/s, Model: BNO055, Bosch, Germany) was fastened to the bottom of the object to measure the roll of the object defined as the angle between the gravitational vector and the vertical axis of the manipulandum, measured in the frontal plane. Object lift onset was obtained as the instant at which either of the lift switches placed under each end of the manipulandum’s base was released by the upward movement of the object from the table’s surface. The sensors’ locations relative to the graspable surfaces were occluded from the participant’s view with a cover plate to prevent visual cues from influencing their fingertip placement. Two arrays of four LEDs (red, ∼5mm) each, mounted on both the inner edges of the cover plate, with the right array placed 10 mm above the left array (relative to the participant), served as cues for finger placement. The width of each of the cues (arrays) was approximately 22 mm. The cover plate was lengthwise perforated (hole diameter ∼ 0.8 mm) along both edges to allow only the light from the LED array to be visible to the participant without revealing their actual position. This served to not divulge any cues about the LED array’s location when switched off. The centre-to-centre distance between the right and left arrays of 10 mm was approximately equal to the average vertical distance between the thumb and the index fingertip, as reported by *Fu et al.* in 2010 (Fu et al., 2010). The non-collinearity of the targets was intended to assist the participants in producing the compensatory moment and diminish fatigue over the course of the experiment. A 400 g mass was placed in the right (relative to the participant) compartment at the base of the manipulandum for all the lifts and was hidden from view to prevent the participants from anticipating the object’s mass distribution. The added mass created an external clockwise moment of 223 N-mm in the frontal plane relative to the participant, and the total weight of the manipulandum was 810g. During the *mixed* block, a second inertial measurement unit (IMU; BNO055, Bosch) was attached to the participants’ wrists to detect movement onset.

### Experimental task and procedure

The experiment was divided into three blocks: unconstrained *(uncon*), constrained (*con*), and *mixed*. The *con* block consisted of fifteen trials. In this block, the participant was presented with visual cues for finger placement throughout the trial. The *uncon* block also had fifteen trials wherein the participants had no visual cues and consequently had the choice of finger placement on the manipulandum. The *mixed* block was created by interspersing a set of *catch* trials within a set of constrained trials. There were fifty trials in this block, among which there were thirty-five constrained trials (referred to across the manuscript as *test* trials), and the rest were *catch* trials. The *catch* trials sat within the *test* trials according to a pseudo-random binary sequence (Fig. 2).

Participants were asked to wash and clean their hands with soap and towel-dry and sit comfortably on a wooden chair. The experiment began with the participants being asked to sit facing the manipulandum with their elbows flexed at ∼ 70^0^ in a parasagittal plane, to align their right shoulder with the midpoint of the manipulandum, and to place their hand (palm facing downward) on a mark (home position) located ∼ 40 cm from the manipulandum. All the trials across all the blocks (*uncon, con*, and *mixed*) had the participants rest their palms in the home position at the beginning of the trial as they heard an auditory cue. For the *con* block and the *mixed* block, the LED arrays intended as visual cues for finger placement were turned on simultaneously. For the *uncon* block, no visual cues were presented at the outset of the trial (Fig. 2). A second auditory cue was presented five seconds after the first tone in all the blocks and signalled the participants to start reaching for the manipulandum with their right hand. The five-second delay was added to amplify the duration of movement planning.

**Fig. 1.**
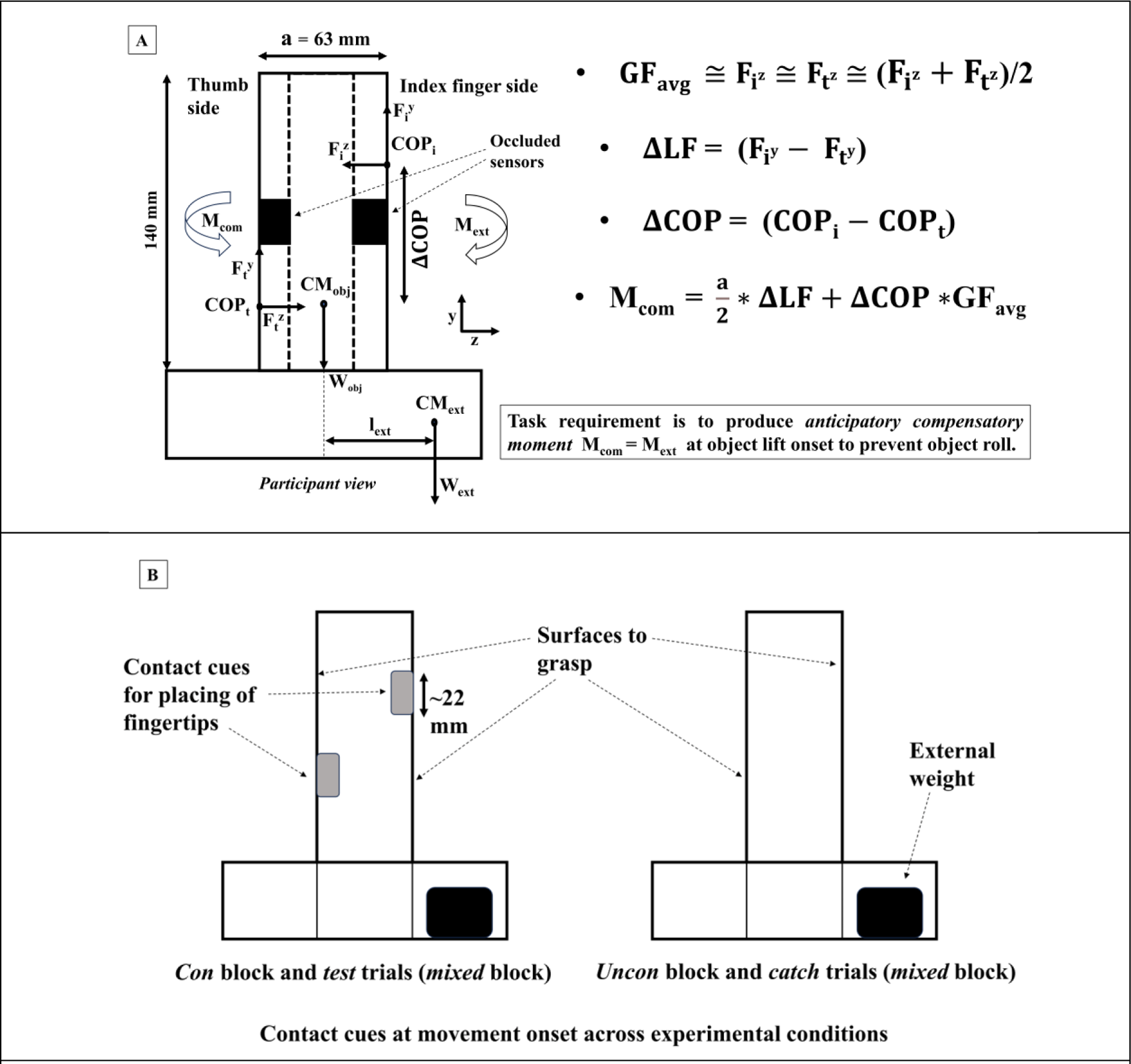
Experimental setup. A. Free-body diagram of the manipulandum showing the fingertip forces exerted by the participants in the normal (F_z_), z direction (grip) and tangential (F_y_), y direction (load) and their respective centres of pressures (COP), the grasp aperture (a) the centre of mass (CM_obj_) and the weight of the object (W_obj_) without considering the external weight, the external weight (W_ext_), and the centre of mass of the added weight (CM_ext_). Participants needed to exert a compensatory moment (M_com_) to balance the external moment (M_ext_ = 223 N.mm) caused by the external weight of 400 gms inserted into the rightmost slot at a distance (l_ext_ = 57 mm) from the centre of the object. The static equilibrium analyses are restricted to the frontal, i.e., the YZ plane. B. Contact cues for all the experimental conditions. The left panel shows the layout of the visual cues (LED array) for the *con* and the *test* trials of the *mixed* block, while the right panel shows the contact condition for the *uncon* and the *catch* trials of the *mixed* block.

Subsequently, as instructed, the participants had to reach for the manipulandum and grasp at the grasp surfaces with the fingertips of the thumb and the index finger, lift it to a comfortable height of (∼12 cm) as indicated by a marker placed beside the object, in an unapprehensive and natural manner, while preventing its tilt on any side, to the extent possible. For the *uncon* block, participants were directed to position their fingertips freely on the grasp surfaces without touching the edges. For the *con* block and the *test* trials of the *mixed* block, they were asked to lift the device by grasping at the visually indicated contact points. For the *catch* trials of the *mixed* block, as soon as the participants heard the second auditory tone and began reaching, the visual cues were switched off by detecting movement onset. The participants were informed about this possibility at the beginning of the *mixed* block. The *catch* trials interspersed within the *test* trials served to reinforce this expectation over the course of the *mixed* block. There was no explicit instruction to the participants regarding where to grasp the object in the *catch* trials after the visual cues were turned off at movement onset. The locations along the graspable panels used to cue the grasp points visually were identical throughout the experiment.

A successful performance required the participants to exert a compensatory moment (M_com_) of the same magnitude but in the opposite direction of the external moment (M_ext_ = 223 N.mm) in an anticipatory manner, i.e., at object lift onset. It needs to be underscored again that since in the *catch* trials, the visual cues were turned off at movement onset as opposed to the *test* trials wherein the visual feedback persisted throughout the trial, the *test* trials essentially were identical to the *con* block except for the uncertainty during the pre-movement planning phase in the former. To get acquainted with placing their fingertips on the cues and the protocol in general, the participants performed two practice trials each for the *uncon*, the *con*, and the *catch* conditions with the external weight in the centre compartment. The three blocks (*uncon*, *con*, and *mixed*) constituting the experiment were counterbalanced, resulting in eighty object lifts in total. Between each block, participants had the chance to rest for a minimum of 5 minutes to minimise fatigue-induced interference in task performance and motor behaviour.

**Fig. 2.**
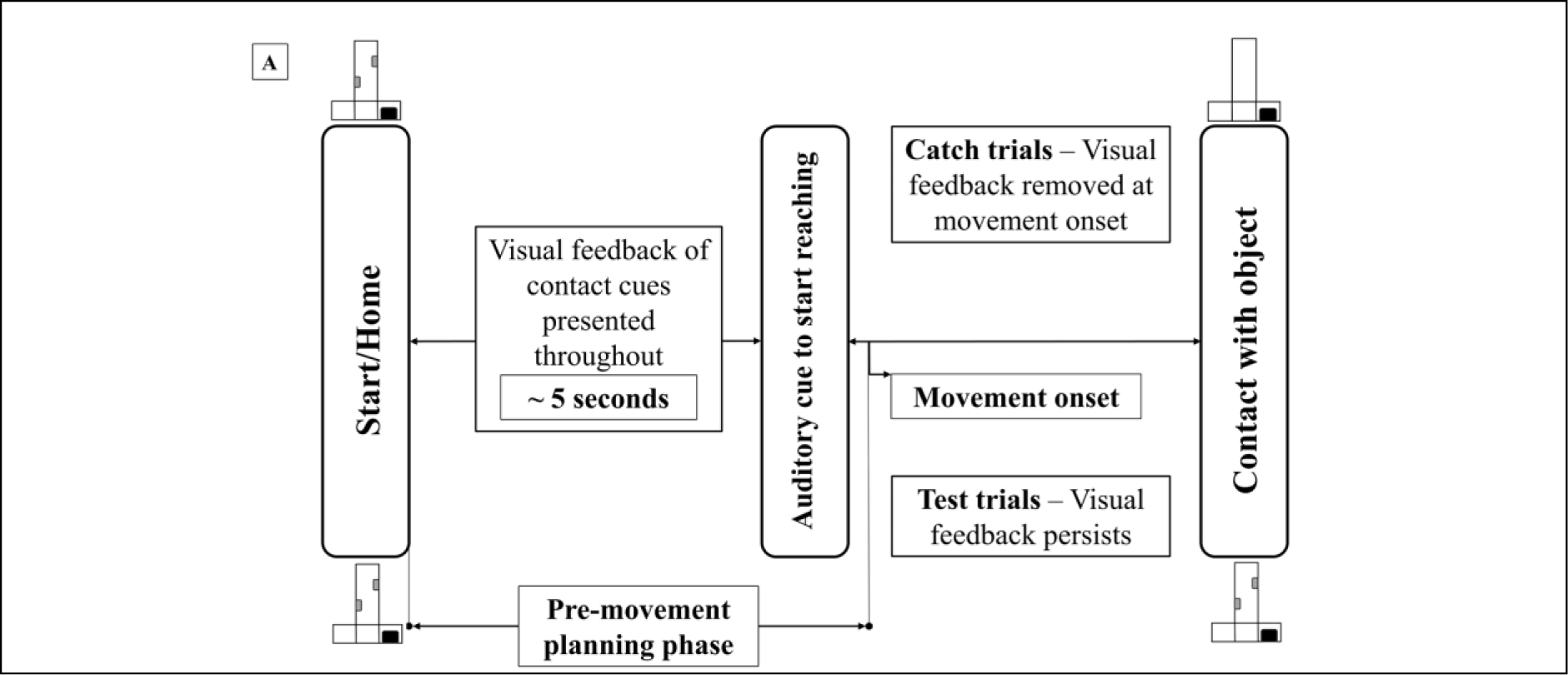
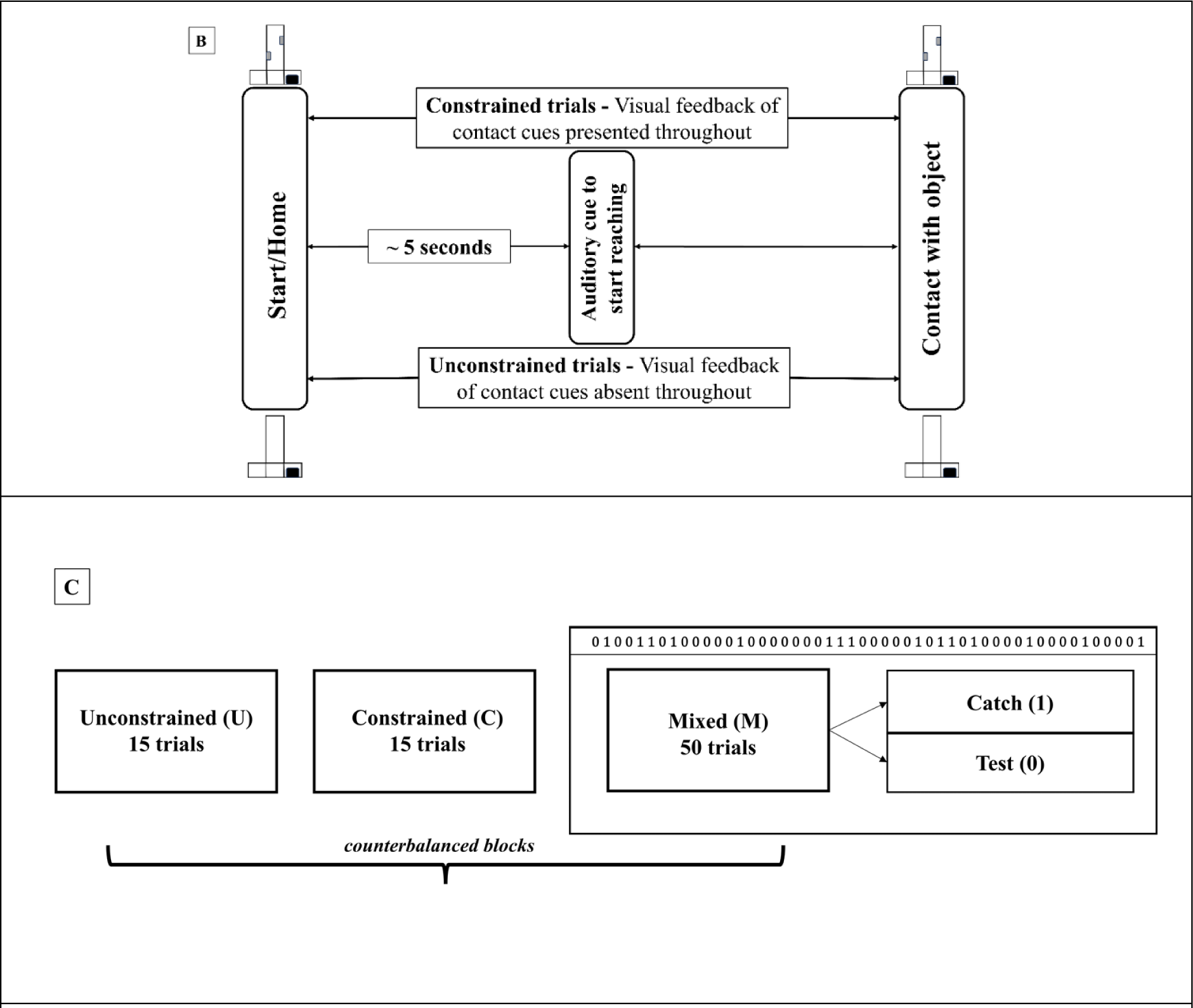
Schematic of the experimental protocol. A. Experimental design for the *mixed* block (*test* and *catch* conditions). B. Experimental design for the constrained (*con)* and the unconstrained (*uncon)* block. C. Binary sequence in the *mixed* block according to which *catch* trials (1) are interspersed within the *test* trials (0). All the conditions, *con, uncon,* and *mixed,* are counterbalanced across participants.

### Data acquisition

The force/moment (F/T) data, the data from the IMUs, and the data from the lift switches were collected using a custom-written LabVIEW code at a sampling rate of 100 Hz. The data were then digitised using a NI DAQ 16-bit A/D board (USB 6225, National Instruments, Austin, TX, USA) and smoothed by running through a second order zero phase lag Butterworth low pass filter (cut-off frequency: 15 Hz). The data from the IMU attached to the object and the IMU attached to the wrist (during the *mixed* block) were digitised through two microcontrollers (Teensy 4.0) and synchronised with the F/T data. A NI DAQ 16-bit A/D board (USB 6002, National Instruments, Austin, TX, USA) was used to acquire data from the lift switches as well as to power and control the LED arrays.

### Data analysis

All the data were analysed offline using MATLAB (Version R2022a, MathWorks, USA). The analyses were based on a few crucial epochs, which are as follows:

*Loading phase* was defined as the duration between the effector’s early contact and object lift onset.

*Early contact* was defined as the instant when the sum of thumb and index grip forces crossed a threshold of 0.1 N and continued to be above it for the next 200 ms.

*Object lift onset* was defined as the instant when either lift switch was released and remained open for the following 50 ms. This is the decisive instant when sensory information about the object dynamics becomes entirely available to the participant.

*Movement onset* was defined as the time when the horizontal acceleration of the hand crossed a threshold ∼ mean + 4 SD of the baseline (Wing & Lederman, 1998).

The *centre of pressure*, i.e., the point of application of the resultant fingertip forces on the grasp surfaces, was calculated from the force and moment components measured from the respective F/T sensors relative to their frames of reference, using the following formula:

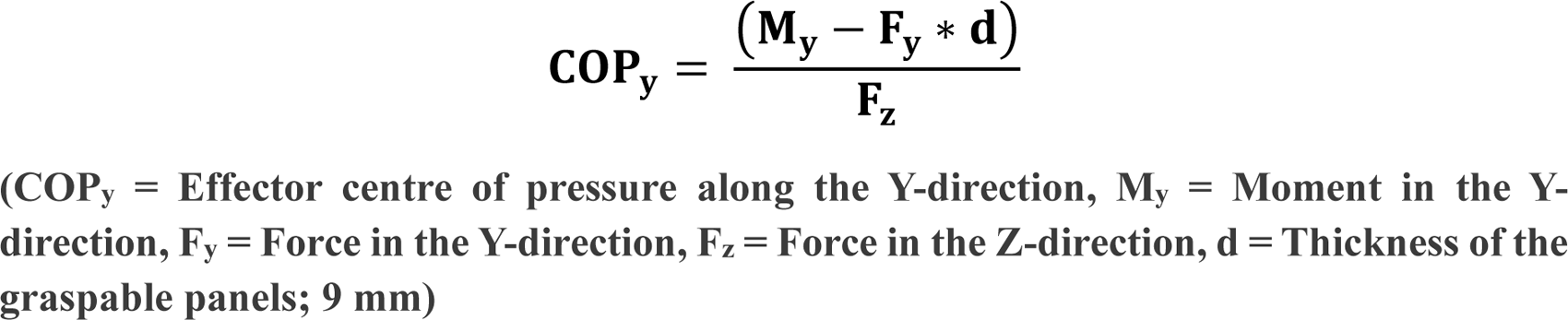

The fingertip forces and the centres of pressure from each of the sensors were used to compute the following performance variables contributing to the compensatory moment:

a. The average of the fingertip grip force values (**GF_avg_**). The grip forces of each side can be reasonably approximated to be equal, i.e., 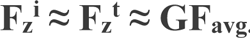
b. The difference between the load forces exerted on the index finger side and the thumb side of the manipulandum, respectively (**ΔLF**), and
c. The vertical distance between the COP on the manipulandum’s index finger and thumb sides, respectively (**ΔCOP**).

The compensatory moment measured at object lift onset (**M_com_**) in the frontal (YZ) plane has been considered an established sign of predictive processes in motor control literature in that it indicates anticipatory force control before complete information through sensory feedback about the object weight and weight distribution becomes available (Fu et al., 2010; Salimi et al., 2000). This compensatory moment for minimising peak object roll can be simplified as a function of the variables mentioned above according to the following equation of static equilibrium, i.e., balancing the eccentricity of the manipulandum caused by the external mass (Fig. 1).

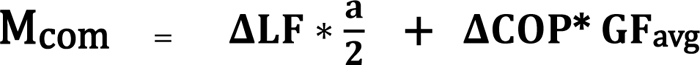

Another variable, Var_ΔCOP (variability in the vertical gap between the fingertip COPs), which is also crucial to the context, was computed for each participant as the standard deviation (SD) of the variable ΔCOP within each experimental condition. All these above performance variables were calculated at object lift onset. As expressed above, the individual terms involved in the expression for the compensatory moment could be broken into the sum of two moments viz.

1. **ΔLF *a/2**: The moment caused by the product of the difference between the index and the thumb tangential forces and half of the grasp aperture (a = 63 mm).
2. **GF_avg_* ΔCOP**: The moment caused by the product of the average grip force and the difference between index and thumb centres of pressure on the grasp panels.

The sign of the exerted moments on the manipulandum is considered positive when directed opposite to the external moment. Hence, from the participant’s viewpoint, moments exerted in the anti-clockwise direction are seen as positive moments and vice versa.

### Spectral analysis of the grip force rate

Grip force rate (GFR) was obtained as the first derivative of the grip force with respect to time. The five-point stencil method was implemented in MATLAB to obtain the derivative. Grip force was defined as the normal component of the fingertip force exerted at the fingertip centre of pressure (COP) on the graspable surfaces. To extract the spectral characteristics of the signal, we used the Continuous Wavelet Transform (CWT) on the GFR signal during the loading phase to quantify the relative intensity of feedback-based corrections in the fingertip force along the lines of *Mojtahedi et al.* (Mojtahedi et al., 2015). The CWT was calculated by integrating the GFR over the duration of the loading phase after multiplying by the scaled and translated versions of the Mexican Hat or the Ricker wavelet function, which is the negative normalised second derivative of a Gaussian function (see *supplementary material* for details).

This analysis aimed to identify bell-shaped signatures in the grip force rate. In motor control literature, bell-shaped profiles of variables such as the wrist velocity during point-to-point fast-reaching movements or the grip force rates while lifting familiar objects are typically thought to indicate feedforward control processes. On the contrary, predominantly feedback-based responses, which are characteristic in sudden target changes during centre-out reaching tasks or when the object weight changes without the knowledge of the participant, exhibit irregular trajectories of motor variables (Ghez & Gordon, 1987; A. M. Gordon et al., 1993; J. Gordon et al., 1995; Jeannerod, 1984; Johansson & Westling, 1988). For teasing out the corrective responses in the grip force rate during the loading phase of the object lift, the Mexican hat function was selected for the CWT analysis because of its resemblance to a bell shape. A metric R_avg_ was computed from the resulting CWT coefficients by varying the scale (pseudo-frequency) and translation parameters and was defined as the ratio of slow bell-shaped components to the sum of slow and fast bell-shaped components as devised in literature (Mojtahedi et al., 2015). The slow bell-shaped component was obtained from a select set of lower pseudo-frequencies (2.08 to 3.125 Hz), while the fast bell-shaped component was obtained from another set of higher pseudo-frequencies (9.09 to 14.28 Hz). In the context of this CWT analysis, it is reasonable to accept that the slow bell-shaped part would quantify the low-frequency feedforward component of force modulation. In contrast, the fast bell-shaped component is believed to identify and represent the high-frequency, feedback-driven adjustments in force. As a result, when we observe larger or smaller values for the R_avg_ metric, it indicates a higher or lower degree of similarity between the grip force rate (GFR) profile and a bell-shaped profile. This, in turn, suggests whether grip force control is predominantly driven by anticipatory (feedforward) or feedback-based mechanisms.

### Statistical analyses

All statistical analyses were performed in the R environment for statistical computing (R Core Team, 2021). Only the last twelve trials of the *con* and *uncon* block were considered for analysis since they represented the learnt trials. Accordingly, the thirty-three *test* trials after the first three trials were used for analysis in the *mixed* block. Throughout the manuscript, results concerning the *mixed* block/condition shall refer to the *test* trials unless otherwise mentioned. We performed a one-way repeated measures ANOVA with the three experimental conditions as the within-participant factors (*con*, *uncon*, and *mixed*) on the R_avg_ of the thumb GFR during the loading phase. For the performance variables measured at object lift onset, viz. M_com_, ΔCOP, GF_avg_, ΔLF, Var_ΔCOP, five separate one-way repeated measures ANOVA with the three experimental conditions as within-participant factors (*con*, *uncon*, and *mixed*) were performed. We also performed linear regression analysis to probe the correlation between ΔCOP and ΔLF across the learnt trials within each experimental condition. For computing the Pearson’s correlation coefficients, the ΔCOP and ΔLF within each condition were normalised by subtracting the average of all trials from the value of each trial and dividing the result by the standard deviation of the trials. Sphericity was checked for the data, and the number of degrees of freedom was adjusted by the Huynh–Feldt (H–F) criterion wherever the assumption was violated. Pairwise post-hoc Tukey tests were performed to examine the significance within conditions. Partial eta-squared (η^2^) was reported as the effect size. Further, the statistical equivalence was tested using the two one-sided t-tests (TOST) (Lakens, 2017) approach for a desired statistical power of 95%. The smallest effect size of interest (SESOI) was chosen as the equivalence bounds. All data in the text is presented as the mean ± SEM. The significance level was set at 0.05 for all the statistical inferences.

## Results

### Task performance

In the present study, we investigated how uncertainty about the grasp context during the pre-movement planning phase affects sensorimotor integration when lifting an object with asymmetrical weight distribution. Despite the differences in grasp context, we expected the participants to learn to anticipate the compensatory moment (M_com_) equally well across all three experimental conditions: *con*, *uncon*, and *mixed*.

**Table 1.** Variables of interest, vis-à-vis the static equilibrium of the manipulandum, and the R_avg_. [CWT-based metric indicating the proportion of slow bell-shaped component in the grip force rate (GFR) spectral composition] **represented as mean ± SEM.** (COP, centre of pressure)

| Variable | *Con* | *Uncon* | *Mixed* |
| --- | --- | --- | --- |
| Compensatory moment ($M_{com}$), N.mm | 164.62 ± 8.97 | 175.19 ± 11.53 | 159.93 ± 6.83 |
| Vertical distance between the index and thumb COPs ($\Delta COP$), mm | 7.82 ± 0.61 | 8.21 ± 1.45 | 8.37 ± 0.58 |
| Average of the fingertip grip forces ($GF_{avg}$), N | 11.58 ± 0.61 | 10.62 ± 0.54 | 10.55 ± 0.61 |
| Difference between the index and thumb load force ($\Delta LF$), mm | 2.14 ± 0.49 | 3.04 ± 0.51 | 2.27 ± 0.32 |
| Variability in the vertical spacing between the fingertip COPs ($Var\_ \Delta COP$), mm | 2.48 ± 0.18 | 4.56 ± 0.62 | 2.68 ± 0.13 |
| $R_{avg}$ of CWT of the thumb GFR ($R_{avg}$) | 0.69 ± 0.02 | 0.62 ± 0.02 | 0.62 ± 0.02 |

The results indeed aligned with our expectations. Since *Fu et al.* (Fu et al., 2010) showed a strong linear correlation between the compensatory moment and peak object roll, which was also present in our results, we interpret task success only through the variable M_com_. All the participants successfully completed the task of lifting the asymmetrically loaded manipulandum while maintaining its vertical alignment. The acquisition of a stable internal representation of the object dynamics is evident from the statistical non-difference of M_com_ measured at lift onset across all experimental conditions (F_1.88,43.41_ = 1.37, p = 0.27, η^2^= 0.056) (see *supplementary material* for analyses on the other performance variables).

### Fingertip force modulation based on the feedback of fingertip position

To probe how participants modulated their fingertip forces depending on the fingertip location on the grasp surfaces, linear regression analyses were performed to quantify the relationship between ΔCOP and ΔLF with the data standardised to zero mean and unit SD across each experimental condition. Expectedly and in line with the previous literature, we observed a stronger linear fit between ΔCOP and ΔLF (Fig. 5) for the *uncon* condition (coefficient of determination, adjusted R^2^ = 0.493, p << 0.0001) as compared to that for the *con* condition (coefficient of determination, adjusted R^2^ = 0.144, p << 0.0001) corroborating previous findings (Davare et al., 2019; Fu et al., 2010; Fu & Santello, 2014; Mojtahedi et al., 2015; Parikh et al., 2020) which have shown the presence of position-dependent fingertip force modulation while grasping an eccentrically loaded object with the freedom of selecting fingertip contact points. The functional significance of such anticipatory negative covariation for the *uncon* condition is rooted in the phenomenon defined as’ digit position to force modulation’ (Corbetta & Santello, 2018). The high variability in fingertip placement and the consequent negative covariation between the load forces applied by the thumb and the index finger and the vertical spacing between the two fingers in the unconstrained condition was evident in our results too. Interestingly, linear regression analysis revealed this phenomenon to be present in the *mixed* condition as well (Fig. 5) (coefficient of determination, adjusted R^2^ = 0.279, p << 0.0001). This is rather surprising since the variability in the fingertip COP in the *mixed* condition was found to be equivalent to the constrained condition (Fig. 4), thus theoretically rendering the need for compensation due to high variability in fingertip COP (Fu et al., 2010) inconsequential. The implications of this result concerning the *mixed* block are elucidated in the discussion section to follow.

### Quantification of feedforward vs. feedback-based force control in the grip force

The variable R_avg_ derived from CWT analysis of the GFR during the loading phase has been established as a key indicator of the nature of force control insofar as the differential weighting of feedforward and feedback-based mechanisms is concerned (Davare et al., 2019; Mojtahedi et al., 2015). As described earlier, smaller and larger values of the R_avg_ indicate a more feedforward or feedback-driven nature of grip force control, respectively. A one-way repeated measures ANOVA on the R_avg_ of thumb GFR during the loading phase showed a significant effect (F_1.83,42.09_ = 18.11, p < 0.001, η^2^= 0.44) of the experimental conditions. Post-hoc pairwise Tukey comparisons revealed the R_avg_ for the *con* condition (mean = 0.698, SD = 0.101) to be greater (q(46) = 7.51, p < 0.001) than the R_avg_ for *uncon* (mean = 0.621, SD = 0.099) condition. This result is consistent with the previous literature and confirms the grasp context-dependent asymmetry in the use of feedback and feedforward-based mechanisms.

**Fig. 3.**
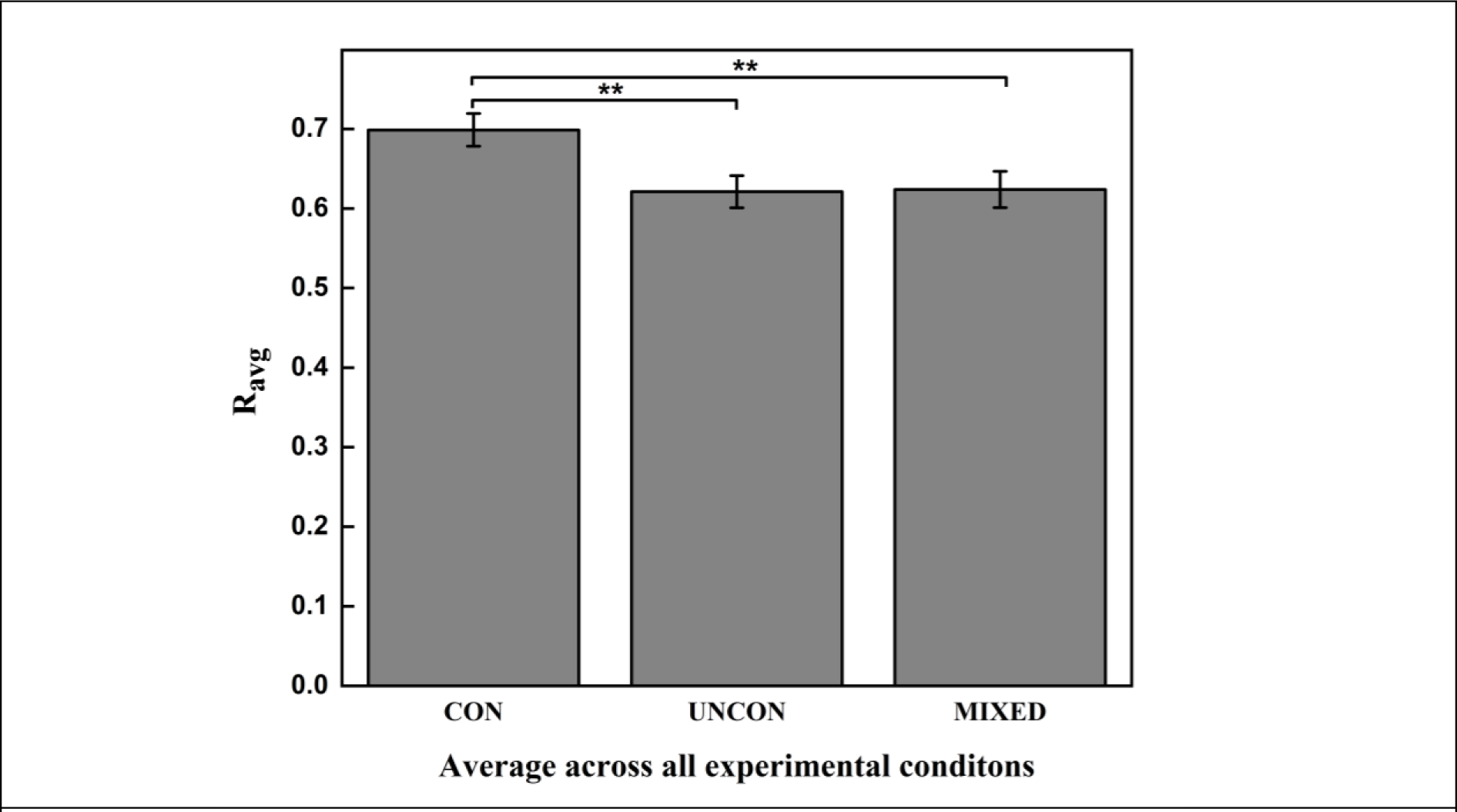
Average of the proportion of slow frequency component (R_avg_) in the spectral composition of the thumb grip force rate (GFR) across all three experimental conditions. The vertical columns and bars indicate the means and the standard errors of means (SE), respectively. Statistical significance is marked by (**) for p < 0.001. The R_avg_ for the *con* condition was found to be statistically greater than both the *uncon* condition and the *mixed* condition. The R_avg_ for the *uncon* and the *mixed* condition were found to be statistically equivalent.

Further post-hoc analyses yielded support for our first hypothesis, in that the R_avg_ for the *con* condition was found to be significantly greater (q(46) = 7.23, p < 0.05) than the R_avg_ for *mixed* condition (mean = 0.623, SD = 0.111). This result is quite remarkable when considering that the *test* trials of the *mixed* block were identical to the *con* trials except for the uncertainty during the period before movement onset. Also, a TOST equivalence test on the R_avg_ between the *uncon* and the *mixed* condition showed statistical significance (t(23) = 4.27, p < 0.0001). Furthermore, to check the variability of the finger contact locations on the grasp panels, a one-way repeated measures ANOVA on Var_ΔCOP showed a significant effect (F_1.14,26.21_ = 11.19, p < 0.0001, η^2^= 0.33) of the experimental conditions. Post-hoc pairwise Tukey comparisons revealed the Var_ΔCOP for the *con* condition (mean = 2.47 mm, SD = 0.86 mm) to be less (q(46) = 6.08, p < 0.0001) than the Var_ΔCOP for *uncon* (mean = 4.56 mm, SD = 3.06 mm) condition. Further, Tukey pairwise comparison between the *uncon* and the *mixed* condition (mean = 2.68 mm, SD = 0.65 mm) showed the variability in the *mixed* condition to be less than that for the *uncon* condition (q(46) = 5.47, p < 0.0001). Also, a TOST test on Var_ΔCOP between the *mixed* and the *con* conditions revealed statistical equivalence (t(23) = 3.09, p < 0.01).

**Fig. 4.**
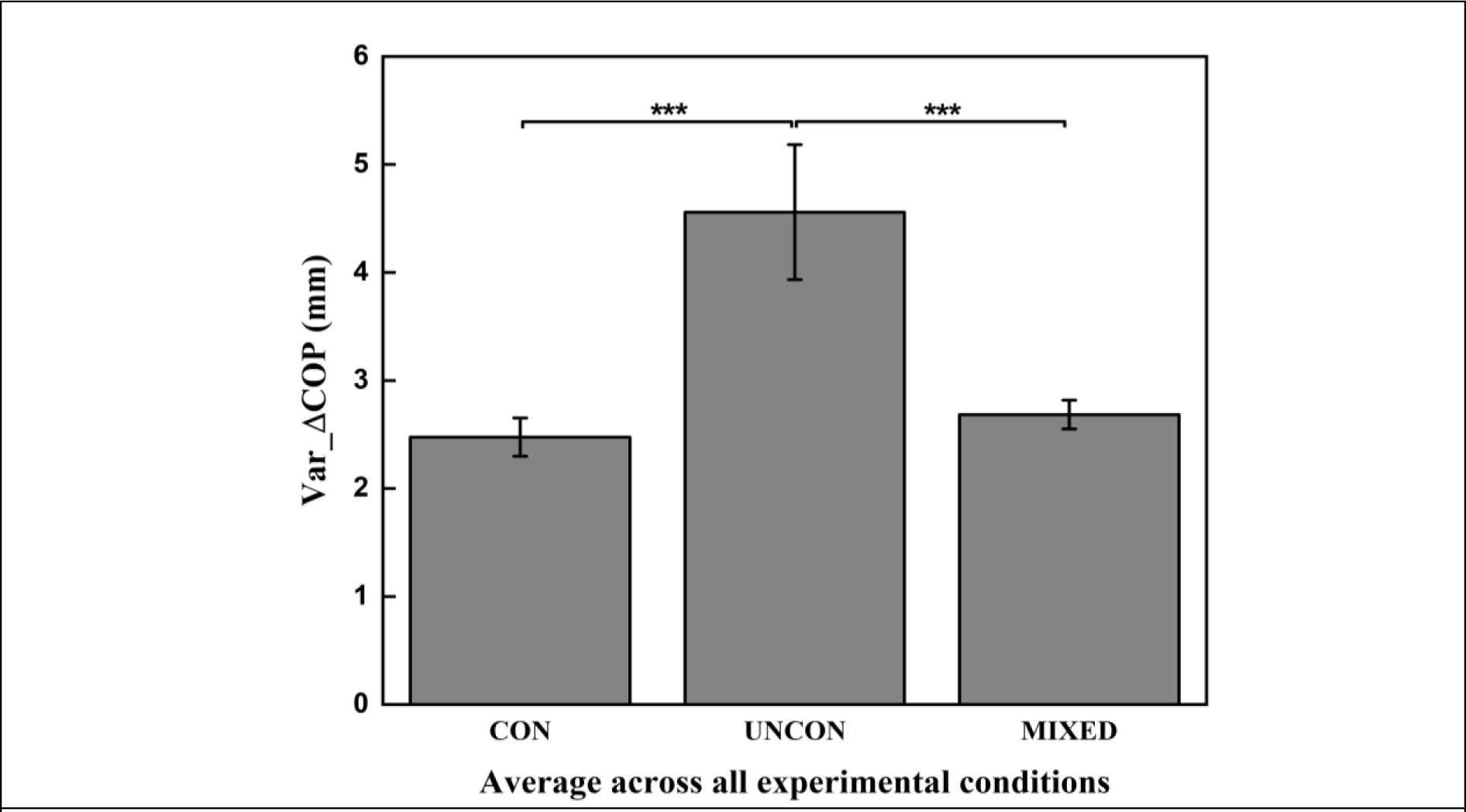
Average of the variability of effector location on the grasp surfaces (Var_ΔCOP) across the three experimental conditions measured at object lift onset. The vertical columns and bars indicate the means and the standard errors of means (SE), respectively. Statistical significance is indicated by (***) for p < 0.0001. The variability for the *uncon* condition was statistically greater than the *con* and the *mixed* condition. The variability for the *mixed* and the *con* conditions were found to be statistically equivalent.

The presence of significantly greater high-frequency feedback-based grip force corrections in the *mixed* block clubbed with the previous result concerning the anti-covariation between ΔCOP and ΔLF points towards a possible deviation from the usage of sensorimotor memories in the *mixed* condition, which considering its task mechanics is somewhat surprising and warrants deep scrutiny. Together, our findings shed new light on how sensorimotor integration underlying anticipatory control is influenced by uncertainty about the grasp context during the planning phase and is expounded in the following section.

**Fig. 5.**
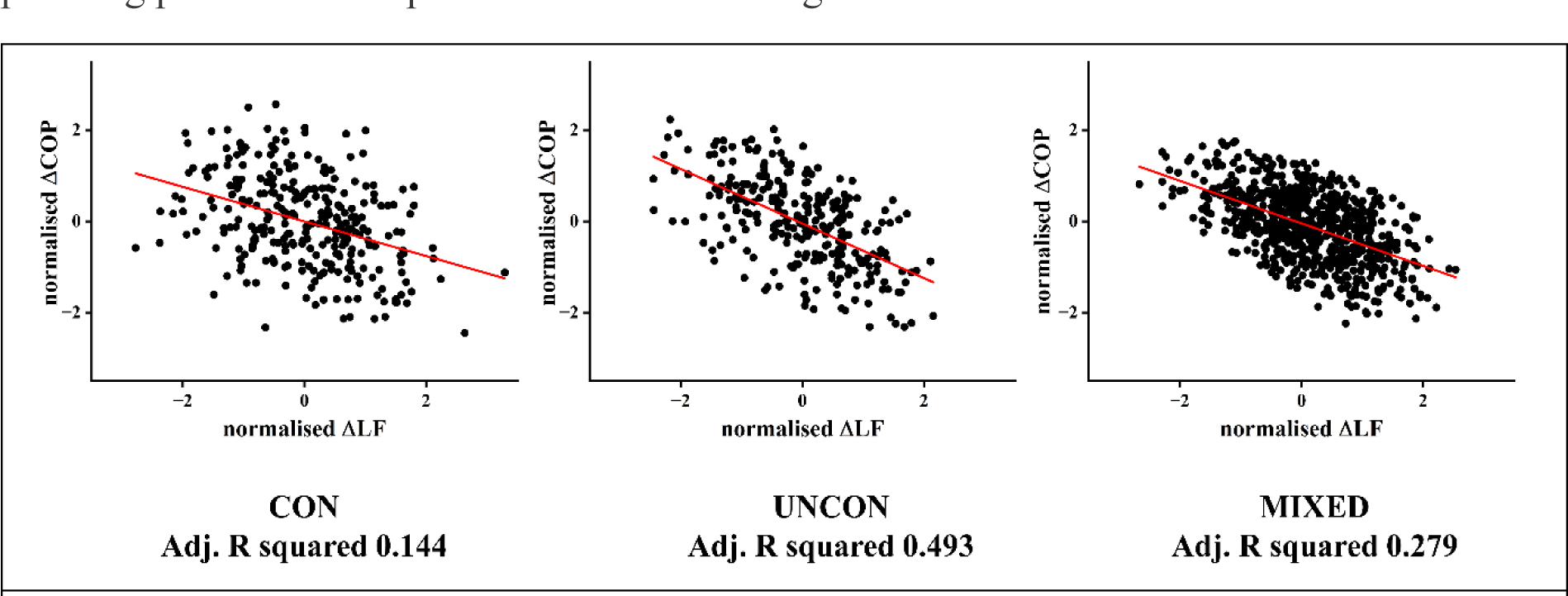
The relationship between ΔCOP and ΔLF is illustrated through scatter plots with the linear regression lines and the corresponding coefficients of determination (R^2^) shown. The data from each condition is from the learnt trials and expressed in the normalised form.

**Fig. 6.**
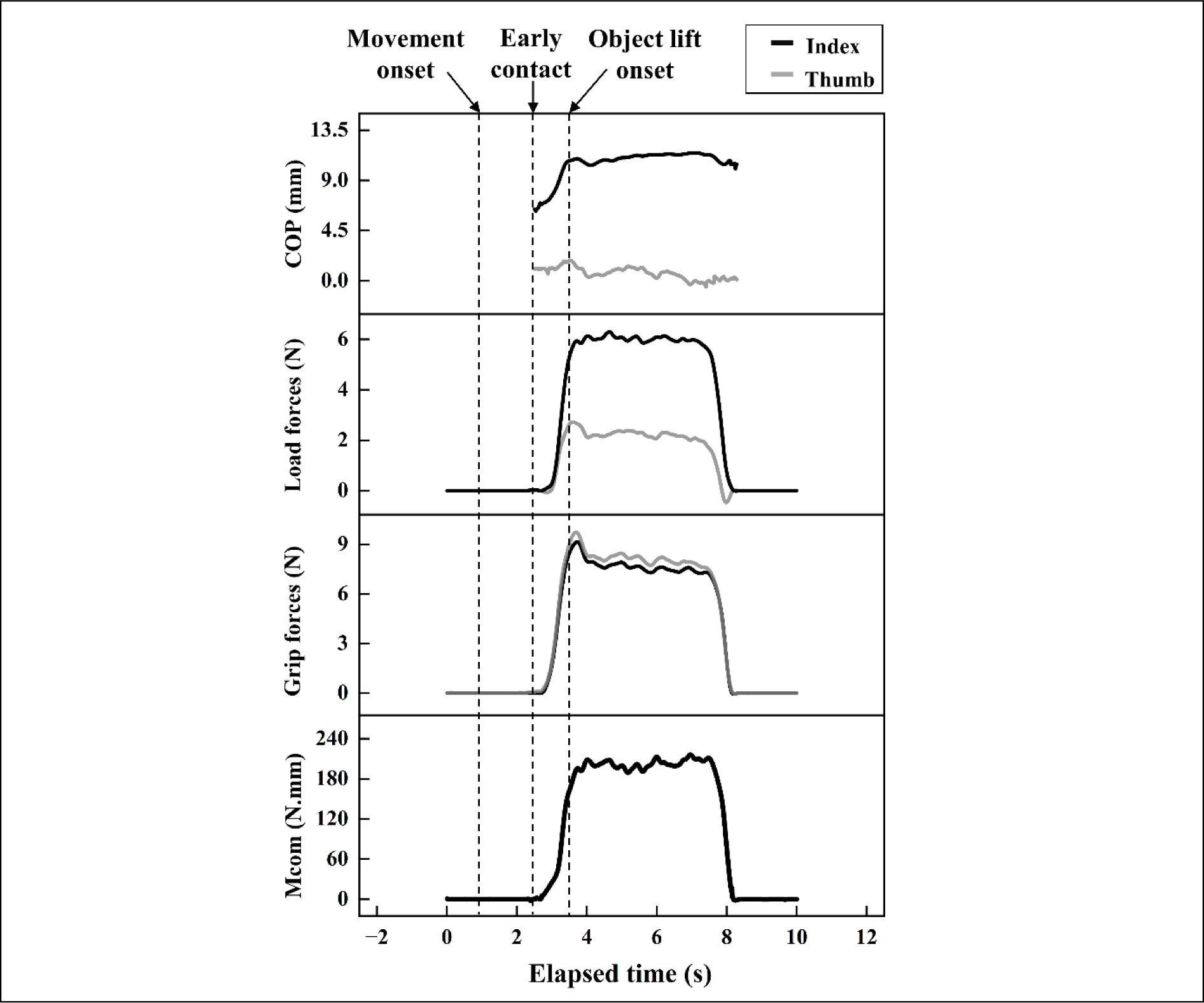
Performance variables contributing to the static equilibrium of the manipulandum are illustrated for one representative trial of the object manipulation task for a *test* trial within the *mixed* block. From bottom to top, the traces denote the compensatory moment (M_com_), grip and load forces of the index fingertip and the thumb and their centres of pressure (COP) on the grasp surfaces. Vertical dotted lines indicate crucial epochs during the considered trial.

## Discussion

The present study employed a unique behavioural paradigm to induce uncertainty about the grasp context during the movement preparation phase, i.e., reaction time before voluntary movement is initiated. Our idea was to examine the role of such uncertainty about the presence of grasp targets in sensorimotor integration vis-à-vis the differential weighting of feedback and feedforward fingertip force control mechanisms in the light of motor planning. The task required the participants to anticipatorily compensate for the external moment in order to lift an object with asymmetrical weight distribution while minimising the roll.

Our results concerning the constrained and the unconstrained conditions replicated previous findings in the literature and substantiate the above framework. Along expected lines, we found an inverse covariation between the load forces applied by the thumb and the index finger and the vertical spacing between the two fingers at lift onset for the unconstrained condition. Such position dependent effector force modulation is primarily not necessary when the position variability is constrained in the manipulandum and was expectedly found absent in our results for the constrained block. A CWT based metric R_avg,_ which was devised to quantify feedback based high frequency (Mojtahedi et al., 2015) corrections in the grip force profile, expectedly in line with previous literature, detected a significant difference between the constrained and the unconstrained conditions in our study.

Interestingly however, we observed support for our hypothesis in that the above measures of effector force control based on the feedback of contact location, viz. firstly, the negative covariation between the load forces applied by the thumb and the index finger(ΔLF) and the vertical spacing between the two fingers (ΔCOP), and secondly, the measure of high-frequency feedback-based corrections (R_avg_), were both found to be significantly higher in the *test* trials of the *mixed* block relative to the constrained trials. This is rather remarkable considering that the only distinguishing feature between *test* trials and the constrained condition was the uncertainty about the grasp context during the pre-movement planning phase in the *mixed* block which resolved at movement onset. This discrepancy between these two behaviourally identical conditions points towards a causal link of the induced uncertainty in anticipatory force coordination during grasping. The following paragraphs are dedicated to probing what we can attribute this anterograde interference to. Feedback-based fingertip force control, which forms the basis for compensating when there is high variability in effector contact points, is rendered largely inconsequential when the fingertip placement cues are constrained or pre-determined. Low fingertip position variability in the *mixed* block relative to the unconstrained condition showed that the participants did not bypass the instructions and, indeed, had used the visual cues to grasp the object. Despite this lack of variability in the fingertip contact points in the *test* trials, as in Fig. 4, our findings report a dominant use of online feedback-based corrective force control over feedforward mechanisms in the *mixed* block. This suggests that in the presence of uncertainty about the grasp context, participants chose to eschew memory-based force control in favour of force modulation reliant on the sensory-haptic feedback of the actual effector location. Why though? One possibility is that the uncertainty before movement initiation hampered retrieval of the previously used sensorimotor map, leading to integrating the fingertip kinematics into the internal representation of the object dynamics on a trial-by-trial basis and computation of a novel motor command each time. However, it is unclear how the retrieval of previously used force maps would be selectively affected and not the ‘high-level’ object representation (Fu et al., 2010) that enabled the re-computation.

It is worthwhile to explore these findings in the light of the neural correlates of motor planning. Motor planning commonly refers to the motor system reaching a favourable preparatory state vis-a-vis voluntary movement. When this preparatory point is attained, and the execution begins, the system’s characteristic dynamics then cause the movement to proceed (Churchland et al., 2010; Shenoy et al., 2013). Recent experiments suggest that motor planning, apart from preparing the motor system for movement, involves changing the neural state of the somatosensory system, likely to facilitate anticipation of the cutaneous signals that emerge as the movement is executed (Ariani et al., 2022; Gale et al., 2021). Additionally, the motor-somatosensory cortical network has been shown to be instrumental in unconstrained object manipulation (Parikh et al., 2020) vis-à-vis integration of online sensory feedback with sensorimotor memories supposedly stored in the M1 (Chouinard, 2006; Jenmalm & Johansson, 1997).

Moreover, uncertainty and probabilistic decision-making have been shown to strongly influence the anterior intraparietal sulcus (aIPS) in the posterior parietal cortex (PPC) (Huettel et al., 2005; Payzan-LeNestour et al., 2013; Soltani & Izquierdo, 2019). The cortex of the intraparietal sulcus has also been implicated extensively in sensorimotor functions such as rapidly building a motor plan during the initial planning stage by incorporating both spatial and pictorial evidence (Verhagen et al., 2012), which is possibly why aIPS lesions can affect seemingly disparate functions like grasping (Binkofski et al., 1998), action organisation (Castiello, 2005), online motor control (Tunik et al., 2005), and tool use (Frey, 2008; Johnson-Frey, 2004). Taken together with the anatomical connectivity of the PPC with both the M1 and the S1 (Edwards et al., 2019), it is plausible that in our study, the uncertainty during the pre-movement planning phase modulated the aIPS of the PPC and consequently affected the preparatory state of the M1-S1 circuitry. We speculate that this anticipatory neural activity caused obligatory sensory processing in the *mixed* condition and led to fingertip force modulation based on the feedback of fingertip location in a situation that did not warrant it.

In conclusion, our study introduces a novel paradigm for investigating the kinetic correlates of uncertainty about the grasp context vis-à-vis self-selection and pre-chosen contact points during object manipulation. We found that such uncertainty during movement preparation anterogradely affects the differential weighting of feedforward and feedback-based fingertip force control in sensorimotor integration. Future studies have been designed to tease out the exact sensorimotor mechanisms at play by probing the neural substrates that could corroborate our behavioural findings.

## Acknowledgements

We thank the Department of Science & Technology, Government of India, for supporting this work, vide Reference no.s SR/CSRI/97/2014 & DST/CSRI/2017/87 under Cognitive Science Research Initiative (CSRI) (awarded to Varadhan SKM). The funders had no role in study design, data collection and analysis, decision to publish, or preparation of the manuscript.

## Author Contributions Statement

Conceptualisation – SD, VSKM; Methodology – SD, VSKM; Formal Analyses – SD; Original Draft – SD; Review and Editing – SD, VSKM.

## Data Availability

We plan on publishing a data descriptor article along with this manuscript. Hence, the data will be made available in due course of time. The data and materials supporting the analyses presented in the paper are available upon reasonable request.

