## Supplemental data for "Grasp context-dependent uncertainty alters the relative contribution of anticipatory and feedback-based mechanisms in object manipulation"

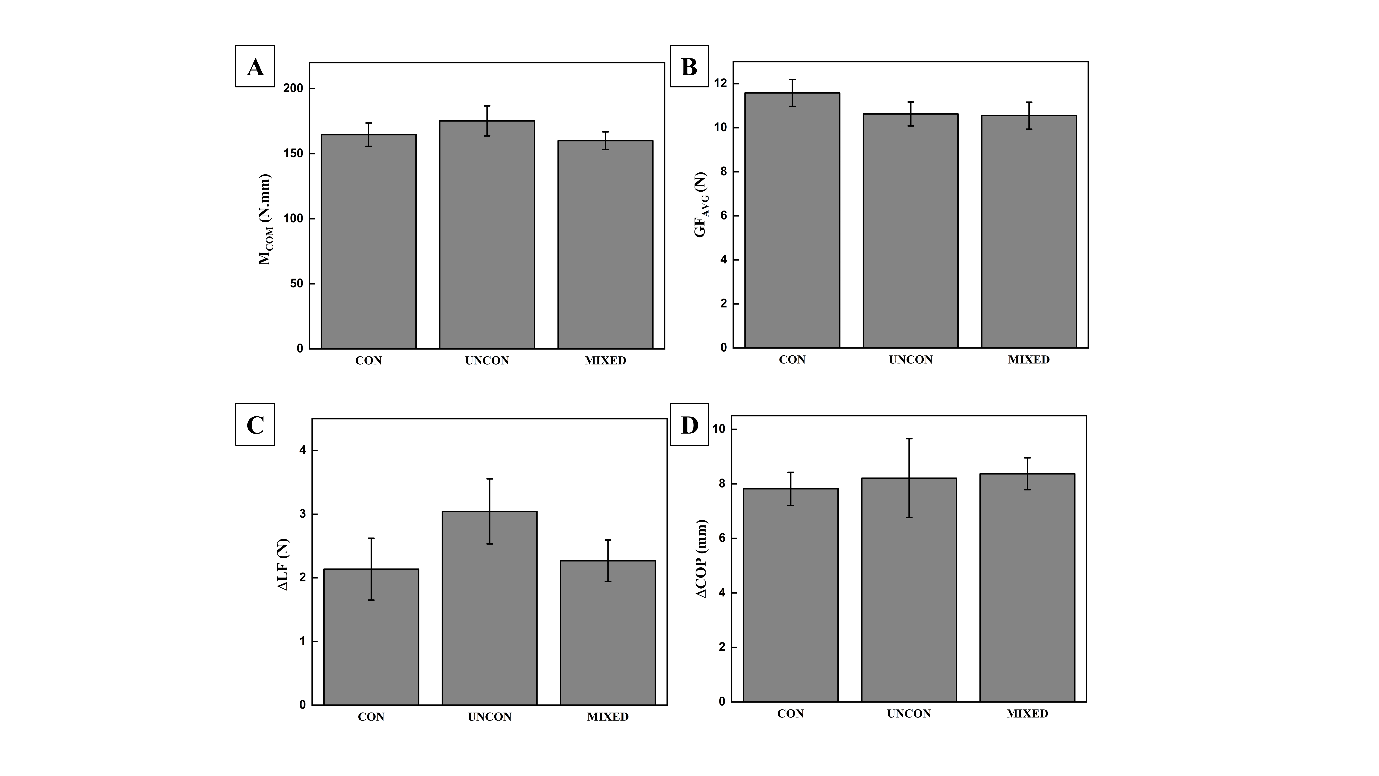

**Fig. S1.** Average of the A. compensatory moment (**M**_com_), B. average of the thumb and index grip forces ($\mathbf{GF}$_avg_), C. difference between the index and thumb load force ($\boldsymbol{\Delta LF}$) D. vertical distance between the index and thumb COPs ($\boldsymbol{\Delta COP}$). All variables were measured at object lift onset. All variables exhibited statistical non difference across all conditions. Vertical bars indicate standard error of means.

1. **Task performance**

Our task required the participants to reach and lift an instrumented manipulandum without tilting it. A successful performance required the participants to exert a compensatory moment (**M_com_**) of the same magnitude but in the opposite direction of the external moment (M_ext_ = 223 N.mm) in an anticipatory manner, i.e., at object lift onset. This compensatory moment for minimising peak object roll can be simplified according to the following equation of static equilibrium:

**M_com_**	_=_	$\boldsymbol{\Delta LF*}\frac{\mathbf{a}}{\mathbf{2}}$	**+**	$\boldsymbol{\Delta COP}$***** $\mathbf{GF}$**_avg_**

This section illustrates task performance results through the relevant variables A. compensatory moment (**M**_com_), B. average of the thumb and index grip forces ($\mathbf{GF}$_avg_), C. difference between the index and thumb load force ($\boldsymbol{\Delta LF}$) D. vertical distance between the index and thumb COPs ($\boldsymbol{\Delta COP}$). All the participants successfully completed the task equally well as is indicated by the statistical non difference of **M_com_** measured at object lift onset (F_1.88,43.41_ = 1.37, p = 0.27, $\eta^{2}$= 0.06). Existing findings regarding the other behavioural measures i.e. $\boldsymbol{\Delta COP} ,\mathbf{GF}$**_avg_** and $\boldsymbol{\Delta LF}$ show similarity between constrained (*con*) and unconstrained (*uncon*) grasping. Our expectation was that the similarity would extend for the *test* trials of the *mixed* condition as well. The results corroborated the literature as well as our expectation in that these measures exhibited statistical non difference across all the three experimental conditions.

- Vertical distance between the index and thumb COPs **(ΔCOP**) : F_1.26,29.15_ = 0.11, p = 0.80, $\eta^{2}$= 0.01
- Average of the fingertip grip forces (**GF_avg_**) : F_2,46_ = 2.03, p = 0.14, $\eta^{2}$= 0.08
- Difference between the index and thumb load force (**ΔLF**) : F_2,46_ = 2.344, p = 0.11, $\eta^{2}$= 0.09

2. **Continuous Wavelet Transform (CWT)**

The Grip force rate (GFR signal of the thumb was used to extract spectral features in the time frequency domain through Continuous Wavelet Transform (CWT) analysis. The continuous wavelet transform (CWT) employs inner products to quantify the resemblance between a signal and a set of analysing functions, where the analysing or basis functions are called wavelets, as denoted as ψ^*^. These wavelets are generated by systematically shifting and either stretching or compressing a ‘mother’ wavelet (ψ), also called scaling. Through a comparison of the signal with the wavelet at different scales and positions, a two-variable function is derived. The CWT coefficients are influenced not only by the values of scale and position but also by the type of the ‘mother’ wavelet. By continuously adjusting the scale parameter *a* and the position parameter *b*, the CWT coefficients, denoted as C(*a,b*), are obtained. If the CWT coefficient for the signal *f(t)* at the location *b* (translation in time) gives a relatively large value, the signal is thought to contain the spectral component corresponding to *a.* For our analysis, we considered the ‘Mexican Hat’ or the ‘Ricker’ wavelet as the ‘mother’ function, which is the negative normalized second derivative of a Gaussian function approximating a bell-shaped profile.

$$\boldsymbol{C}\left( \boldsymbol{a,b} \right)\boldsymbol{=}\int_{\boldsymbol{-\infty}}^{\boldsymbol{\infty}} \boldsymbol{f}\left( \boldsymbol{t} \right)\frac{\boldsymbol{1}}{\boldsymbol{a}}\boldsymbol{\psi}^{\boldsymbol{*}}\left( \frac{\boldsymbol{t-b}}{\boldsymbol{a}} \right)\boldsymbol{ⅆt}$$

The frequency *f* (also called pseudo-frequency) relates to the scale *a* through an inverse proportionality according to the following equation:

$$\boldsymbol{f=}\frac{\boldsymbol{f}_{\boldsymbol{c}}\boldsymbol{\cdot}\boldsymbol{f}_{\boldsymbol{s}}}{\boldsymbol{a}}$$

where $f_{c}$ is the central frequency of the mother Mexican Hat wavelet, and $f_{s}$ is the sampling frequency. Thus, larger scale values correspond to smaller frequencies and vice-versa. For extraction of the derived feature **R_avg_,** we followed the procedure as devised by *Mojtahedi et al.* (Mojtahedi et al., 2015). Briefly, the CWT coefficients corresponding to extracted from the with a select et of 5 slow and 5 fast bell-shaped functions. The slow bell-shaped function is equivalent to the lower frequency (or higher scale) or longer period. Conversely, a fast bell-shape, which means that the Mexican Hat is faster or has a shorter duration, is associated to higher frequency (lower scale) or shorter period. The slow component was defined as the average of five higher scales (lower frequencies) of bell-shaped wavelets and the fast component was defined as the average of five lower scales (higher frequencies) of the bell-shaped wavelets. To incorporate information about both the slow and the fast component, a metric **R_avg_** was obtained by averaging the ratio of the slow component to the sum of the slow component and the fast component *R(b)* over the loading phase duration. The table below shows the scales associated with the different pseudo frequencies, Considering the cut off frequency of 15 Hz for the filter, the highest pseudo frequency was 14.28 Hz, and the lowest frequency was taken to be 2.08 Hz (Mojtahedi et al., 2015).

**Table S1**. **Pseudo-frequencies associated with the selected set of wavelets with lower and higher scales**

| Lower scales,*a*  (unitless) | Shorter periods (ms) | Higher pseudofrequencies,*f* (Hz) | Higher scales,*a* (unitless) | Longer periods (ms) | Lower pseudofrequencies, *f* (Hz) |
| --- | --- | --- | --- | --- | --- |
| 1.75 | 70 | 14.28 | 8 | 320 | 3.12 |
| 2.00 | 80 | 12.50 | 9 | 360 | 2.77 |
| 2.25 | 90 | 11.11 | 10 | 400 | 2.50 |
| 2.50 | 100 | 10.00 | 11 | 440 | 2.27 |
| 2.75 | 110 | 9.09 | 12 | 480 | 2.08 |

We used the MATLAB function ‘*cwt*’ to obtain the CWT coefficients of the GFR signals. The *cwt* function uses L1 normalisation as opposed to the commonly used L2 normalisation where the normalisation is done by multiplying with $\frac{\mathbf{1}}{\sqrt{\mathbf{a}}}$. This causes the high frequency amplitudes or the lower scale amplitudes to diminish more than the low frequency peaks. Whereas L1 normalisation employs multiplying by $\frac{\mathbf{1}}{\mathbf{a}}$ thereby rendering a more accurate representation of the signal.
